# The RhoGEF protein Plekhg5 regulates medioapical actomyosin dynamics of apical constriction during *Xenopus* gastrulation

**DOI:** 10.1101/2022.08.31.506049

**Authors:** Austin T. Baldwin, Ivan K. Popov, Ray Keller, John B. Wallingford, Chenbei Chang

**Affiliations:** Department of Molecular Biosciences, University of Texas at Austin, Austin, TX 78712; Department of Cell, Developmental and Integrative Biology, University of Alabama at Birmingham, 1720 2^nd^ Avenue S, Birmingham, AL 35294; Biology Department, University of Virginia, Charlottesville, VA 22903

**Keywords:** apical constriction, bottle cells, actomyosin, *plekhg5*, *Xenopus*

## Abstract

Apical constriction results in apical surface reduction in epithelial cells and is a widely used mechanism for epithelial morphogenesis during embryo development. Both medioapical and junctional actomyosin remodeling are involved in apical constriction, but the deployment of medial versus junctional actomyosin in specific developmental processes has not been fully described. Additionally, genetic regulation of actomyosin dynamics during apical constriction is poorly understood in vertebrate systems. In this study, we investigate actomyosin dynamics and their regulation by the RhoGEF protein Plekhg5 in *Xenopus* bottle cells. Using live imaging and quantitative image analysis, we show that bottle cells assume different shapes, with rounding bottle cells constricting earlier in small clusters followed by fusiform bottle cells forming between the clusters. Though both medioapical and junctional actomyosin accumulate as surface area decreases, apical constriction is better correlated with medioapical actomyosin localization, which may promote formation of microvilli in the apical membrane. Knockdown of *plekhg5* disrupts both medioapical actomyosin activity and apical constriction but does not affect initial F-actin dynamics. Taken together, our study reveals distinct cell morphologies, uncovers actomyosin behaviors, and demonstrates the crucial role of a RhoGEF protein in controlling actomyosin dynamics during apical constriction of bottle cells in *Xenopus* gastrulation.

## Introduction

Apical constriction, an active process via which cells contract their apical surface, is one of the driving forces of epithelial morphogenesis (Sawyer *et al*., 2010; Martin and Goldstein, 2014). Cells undergoing apical constriction often elongate their basolateral compartments and exert pulling forces on their neighbors. Apical constriction is used reiteratively during embryonic development to bend and fold epithelial sheets, such as during gastrulation movements in various animal species, folding of the neural ectoderm into a closed neural tube, lumen and tube formation during organogenesis, and sensory placode morphogenesis. Increasing our depth of understanding of the molecular machinery controlling apical constriction can therefore provide broad insight into epithelial remodeling crucial for embryogenesis.

Apical constriction is regulated by dynamic and spatially restricted reorganization of cytoskeleton networks (Martin and Goldstein, 2014). Both actomyosin and microtubules have been implicated in this process (Rogers *et al*., 2004; Lee *et al*., 2007; Lee and Harland, 2007; Booth *et al*., 2014; Ko *et al*., 2019; Le and Chung, 2021), though the roles of actomyosin have been investigated in more depth. Recruited to apical cell-cell junctions by adhesion complexes, filamentous actin (F-actin) and its associated motor protein non-muscle type II myosin form a circumferential junctional belt that facilitates establishment and maintenance of cell adhesion. Contraction of junctional actomyosin can lead to reduction of apical membrane size by a purse-string mechanism. Enhanced junctional actomyosin activity consistent with purse-string constriction has been observed in ectopic apical constriction induced by specific regulators in epithelial cell cultures, in wound healing responses, and in dorsal closure during *Drosophila* development (Bement *et al*., 1993; Nakajima and Tanoue, 2011, 2012; Kamran *et al*., 2017; Kiehart *et al*., 2017; Yano *et al*., 2021).

Recently, studies in multiple systems reveal that apical constriction can also be mediated by assembly of actomyosin in the medial region of the apical cell cortex. The medioapical actomyosin undergoes dynamic remodeling and engages cell junctions to actively shrink the apical cell surface. This mechanism is utilized during *Drosophila* and *C. elegans* gastrulation, lens placode invagination, and in the neural plate cells during neural tube closure (Martin *et al*., 2009; Plageman *et al*., 2011; Roh-Johnson *et al*., 2012; Lang *et al*., 2014; Martin and Goldstein, 2014; Christodoulou and Skourides, 2015; Jodoin *et al*., 2015; Chanet *et al*., 2017; Goldstein and Nance, 2020). A hybrid strategy of employing both junctional and medioapical actomyosin has also been observed during apical constriction of neural ectoderm cells in *Xenopus* during neurulation (Baldwin *et al*., 2022). Thus, apical constriction mechanisms are both varied and complex, and understanding mechanisms employed in particular processes will involve observing both medial and junctional actomyosin simultaneously.

One of the prominent examples of apical constriction is the formation of bottle cells during *Xenopus* gastrulation (Keller, 1981; Hardin and Keller, 1988; Kurth and Hausen, 2000; Kurth, 2005; Lee and Harland, 2007, 2010). Bottle cells reduce their apex, elongate along the apicobasal axis, and expand their basal compartments during gastrulation. Pigment granules concentrate underneath apical cell membrane as apical surface reduces, making the appearance of darkly colored cells the hallmark of apical constriction during *Xenopus* gastrulation. The bottle cells form initially from the dorsal side and spread toward lateral and ventral quadrants to form the blastopore lip. The change in morphology of bottle cells leads to invagination of surface epithelium that coordinates with tissue involution in the underlying mesoderm. When bottle cells are removed surgically or prevented from forming by gene manipulations, *Xenopus* embryos can still complete gastrulation through tissue involution at ectopic sites that buckle due to other morphogenetic movements. However, imprecision of the tissue involution sites often leads to embryonic defects, such as shortening of anterior archenteron and malformation of head structure (Keller, 1981; Hardin and Keller, 1988; Popov *et al*., 2018). This highlights the importance of bottle cells in conferring precision and robustness of gastrulation movements.

Apical constriction of bottle cells is regulated by signals controlling mesendoderm cell fate and polarity, such as those of nodal and planar cell polarity (Kurth and Hausen, 2000; Choi and Sokol, 2009; Ossipova *et al*., 2015). Our recent studies reveal that a RhoGEF gene, *plekhg5*, is transcriptionally activated by nodal signaling and is required for apical constriction induced by ectopic nodal expression in the animal region or in endogenous bottle cells (Popov *et al*., 2018). Plekhg5 stimulates Rho activation and likely regulates actomyosin dynamics via Rho effectors, such as Diaphanous, which facilitates assembly of F-actin bundles, and ROCK, which activates myosin II through phosphorylation of regulatory myosin light chain (MLC, (Mulinari *et al*., 2008; Massarwa *et al*., 2009; Mason *et al*., 2013; Rousso *et al*., 2013)). Analysis of F-actin and pMLC indeed shows that knockdown of *plekhg5* reduces their apical accumulation in presumptive bottle cells and prevents cell shape changes of bottle cells (Popov *et al*., 2018). However, the behaviors of actomyosin in normal bottle cells and how these are altered in *plekhg5* knockdown embryos are not known.

In this study, we tackle the following questions. How is apical actomyosin remodeled during apical constriction of bottle cells? Do junctional, medioapical, or combined actomyosin activities drive apical surface reduction? What are the kinetics of actomyosin accumulation and apical surface reduction? And how does *plekhg5* regulate actomyosin dynamics in bottle cells? Using live cell imaging and quantitative image analysis approaches, we show that medioapical actomyosin plays a crucial role in apical constriction of bottle cells, and *plekhg5* regulates accumulation of F-actin and myosin in the medioapical cortex without affecting initial F-actin dynamics. Our data address an important knowledge gap about cytoskeleton regulation by a RhoGEF protein during apical cell constriction of bottle cells in *Xenopus* gastrulation, an important developmental process in a vertebrate species.

## Results

### Heterogeneous morphology of the bottle cells

Bottle cell morphology was studied originally by scanning electron microscopy (SEM) in both en face and sagittal views, though molecular studies of bottle cell formation tend to rely on the appearance of surface pigmentation and the side views of the bottle cell shape (Hardin and Keller, 1988; Kurth and Hausen, 2000; Lee and Harland, 2007; Choi and Sokol, 2009; Lee and Harland, 2010; Ossipova *et al*., 2015). To examine apical cell morphology during bottle cell formation, we first labeled the cells by expressing an mRNA encoding membrane-mCherry (mem-mCherry) in the dorsal marginal zone cells. The surface of the labeled epithelial cells was examined at early gastrula stages by confocal microscopy. We observed that bottle cells displayed distinct morphologies. Clusters of small round cells (yellow arrow in **Fig. 1A**) were often flanked by fusiform cells that displayed circumferential elongation with reduced animal-vegetal cell length (blue arrow, **Fig. 1A**). The distinct cell shapes implied that bottle cells could be formed by both isotropic and anisotropic apical constriction to adopt different morphologies.

**Figure 1.**
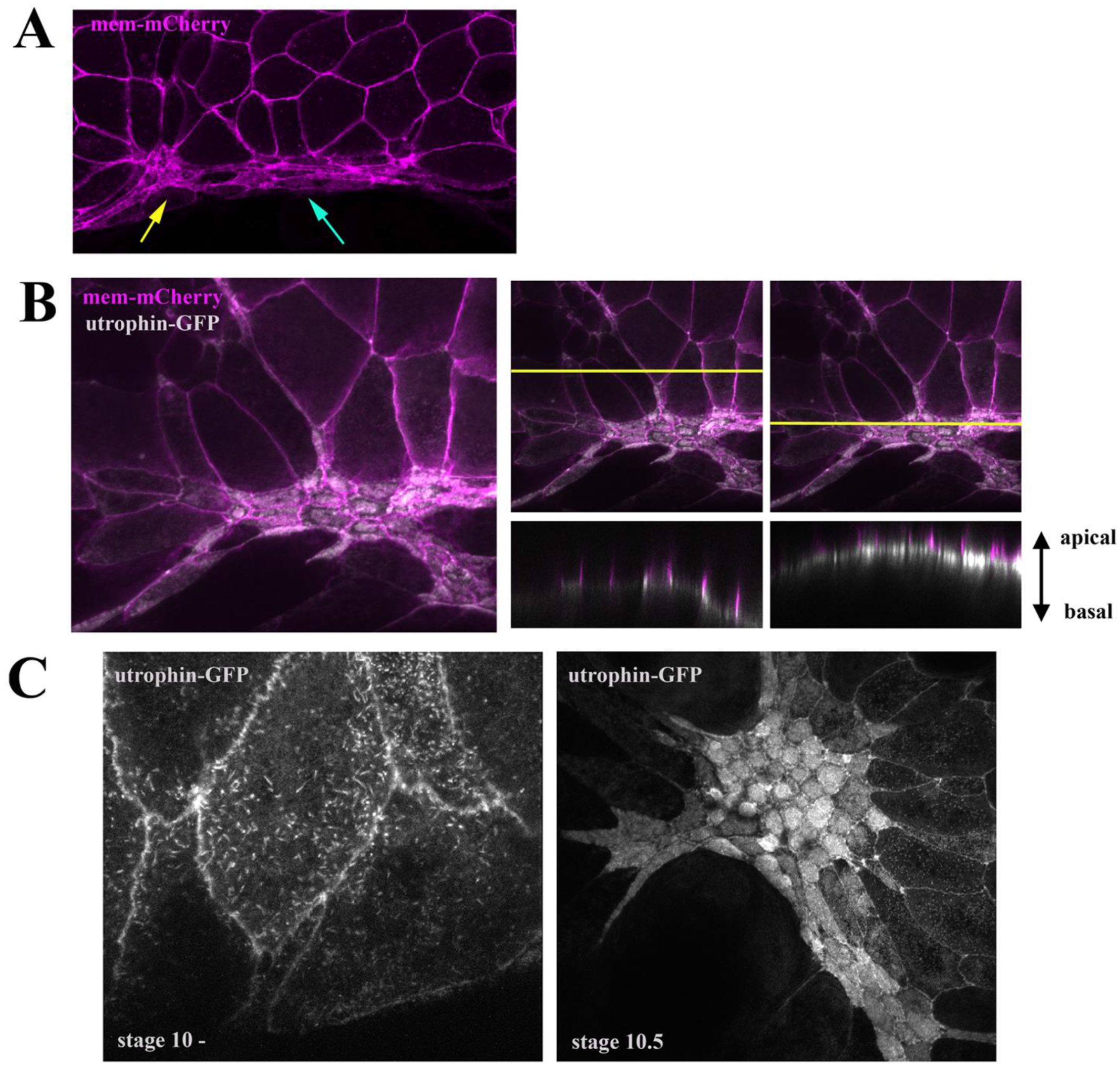
Distinct cell shapes, apical actin accumulation, and microvilli in bottle cells. A) Membrane mCherry – labeled bottle cells show two distinct morphologies, with round cells (yellow arrow) interspersed with fusiform cells (blue arrow). B) Co-expression of membrane mCherry with utrophin-GFP reveals accumulation of F-actin predominantly in the medial regions of apical cell cortex. The right two panels show the z-view of F-actin and mem-mCherry signals in regions without or with apical constriction. C) Formation of microvilli starts prior to substantial reduction of surface area (stage 10, left panel), with dense microvilli seen in both round and fusiform bottle cells at later stages (stage 10.5, right panel).

### Medioapical F-actin distribution in the bottle cells

To investigate whether junctional F-actin belts or medioapical F-actin networks supply the primary constrictive forces in the bottle cells, we co-injected mem-mCherry RNA with that of utrophin-GFP, which encodes the F-actin binding protein Utrophin conjugated with green fluorescent protein. Utrophin-GFP signal was seen primarily at the cell-cell junctions in non-constricting (or not yet constricting) cells but enriched at the apical cell compartment in the bottle cells (**Fig. 1B**). Though junctional F-actin was present in the bottle cells, most of the increased signals were detected in the medioapical region (left panel, **Fig. 1B**). Side views of utrophin-GFP distribution revealed that the signal was not simply accumulated under the apical cell membrane but formed discrete bundles perpendicular to the plasma membrane (right panels, **Fig. 1B**). The patterns appeared similar to the microvilli observed previously on the apical surface of the bottle cells by scanning and transmission electron microscopy studies (Keller, 1981; Hardin and Keller, 1988; Kurth and Hausen, 2000; Lee and Harland, 2010). Close-up confocal images indeed confirmed that F-actin-rich microvilli formed on the surface of emerging bottle cells prior to obvious reduction of their surface areas and became densely packed when apical constriction decreased apical surfaces of the bottle cells (**Fig. 1C**). The results imply that generation of high-density microvilli may be a critical step in reduction of apical area and serve to preserve surface membrane during bottle cell formation.

### Dynamic F-actin remodeling during bottle cell formation

To gain insight into F-actin dynamics during apical constriction of bottle cells, we performed live imaging of embryos co-injected with mRNAs of mem-mCherry and utrophin-GFP (**Suppl. Movies 1 and 2**). Dynamic reorganization of F-actin puncta at the subapical cortex was visible prior to substantial reduction of apical cell surface (arrowheads, top panels, **Fig. 2A**). As the intensity of subapical F-actin was increased with time, the apical cell areas were reduced. The enhanced F-actin signal was seen under the entire medial-apical cell membrane without strong junctional preference (**Suppl. Movies 1 and 2 and Fig. 2**). Tracking F-actin localization at both the medial and junctional domains of the apical surface (**Fig. 2B**) in 132 cells from 3 embryos over the course of apical constriction using Tissue Analyzer revealed a stronger negative correlation between medial F-actin localization and apical area (r = −0.62) (**Fig. 2C**) than junctional F-actin (r = −0.33) (**Fig. 2D**). These results suggest that medial F-actin contractility may be the stronger driver of apical constriction in the bottle cells, similar to the anterior *Xenopus* neural ectoderm (Baldwin *et al*., 2022).

**Figure 2.**
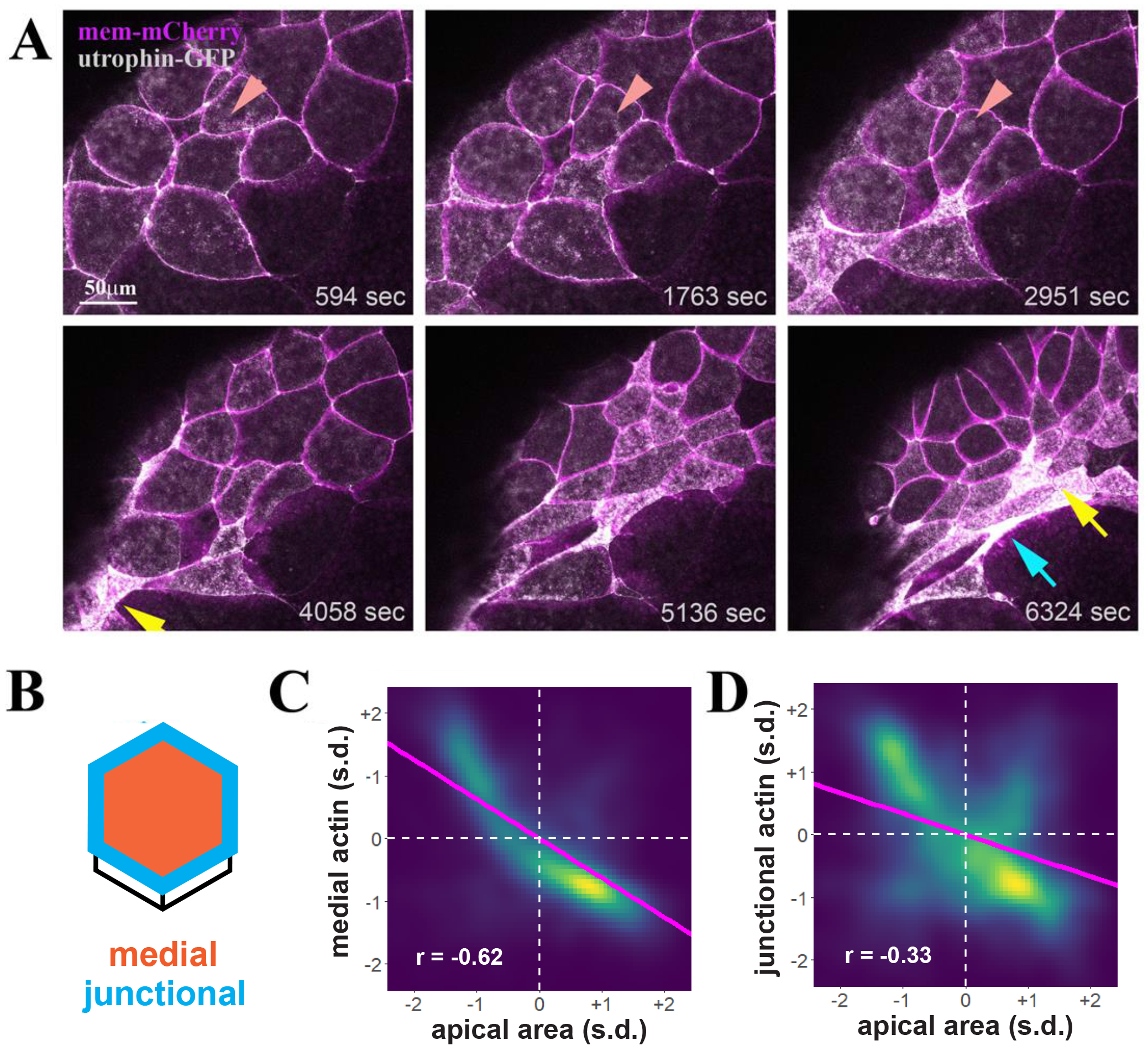
Dynamic F-actin remodeling accompanies apical surface reduction during bottle cell formation. A) Selected frames from a time-lapse movie demonstrate dynamic F-actin organization prior to overt decrease in apical cell area (top panels). The dynamic F-actin signal in a single cell is marked by the arrowheads. As apical constriction commences, groups of cells constrict first and take round shape (yellow arrow), with neighboring, later constricting cells assuming fusiform (blue arrow). B) Schematics of quantification of medial (orange) and junctional (blue) F-actin intensity within single cells. C&D) Density plots of apical cell area vs medial (C) and junctional (D) F-actin. Magenta line indicates linear model. s.d. = standard deviations. n = 11748 observations of 132 cells. C) An inverse correlation of medial F-actin intensity and apical area is uncovered, with the r value of −0.62. D) Junctional F-actin intensity is less correlated with the apical area, with the r value of −0.33.

In generating movies of bottle cell apical constriction, we observed that apical constriction was initiated within clusters of cells spaced some distance apart along the blastopore lip, with cells between the clusters constricting at later times (**Suppl. Movies 1 and 2**). The early constricting cells tended to have a round morphology, reflecting possible isotropic contraction forces (yellow arrows, **Fig. 2A and 3A**). The cells between the clusters adopted a fusiform shape that seemed to be stretched by constricting neighboring cells in the clusters (blue arrows, **Fig. 2A and 3A**; **Suppl. Movies 1 and 2**). Consistent with formation of microvilli in bottle cells, distinct F-actin and membrane mCherry puncta could often be seen at the surface of the constricting cells (**Suppl. Movies 1 and 2**). Quantification of the changes in cell shape using the “stretch” parameter calculated by Tissue Analyzer (Aigouy *et al*., 2010) (**Fig. 3B)** showed that within a cluster of constricting cells, a small number of extremely constrictive cells became rounder (decreased stretch) while surrounding cells took on fusiform shapes (increased stretch), suggesting that apical constriction can occur both anisotropically and isotropically in bottle cells (**Fig. 3C&D**).

**Figure 3.**
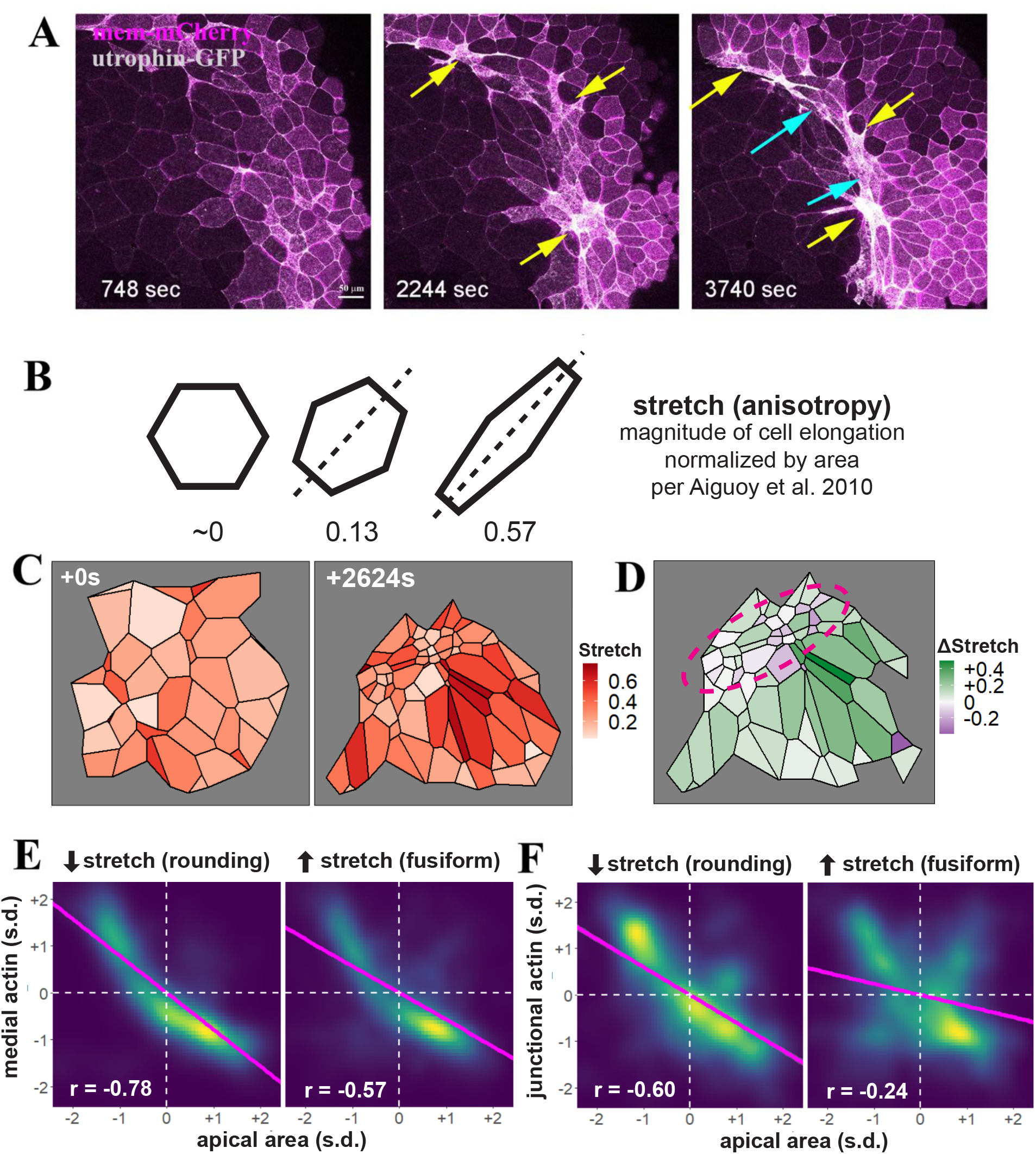
Distinct cell shape changes accompany bottle cell formation. A) Selected frames from a time-lapse movie reveal that apical constriction does not spread from a single point. Clusters of cells some distance apart can constrict around similar time. These cells tend to constrict early and form round shape (yellow arrows), with the cells in between stretching in the circumferential direction to take the fusiform shape (blue arrows). B) Quantification of cell stretch by Tissue Analyzer, whereby the magnitude of cell elongation is normalized by cell area to generate a number ranging from 0 to 1, such that cells with the same shape but different areas will have the same stretch. C) An example of heterogeneous cell constriction and stretch based on quantitative analysis of a time-lapse movie from the first and the last frame. D) Mapping of changes of cell stretch onto the embryo images reveals that round, no-stretching cells are juxtaposed to the cells with increasing stretch in the blastopore lip (purple circle). The neighboring cells animal to the blastopore lip also display distinct changes in stretch, with cells showing strong stretch preferentially abutting the round bottle cells. E&F) Quantification of medial (E) or junctional (F) F-actin intensity reveals a stronger correlation with the round bottle cells than with the fusiform bottle cells. Magenta line indicates linear model. s.d. = standard deviations. n = 3171 observations for cells decreasing stretch (rounding) and 8709 observations for cells increasing stretch (fusiform).

We further explored differences in the contractile dynamics of round versus fusiform cells and found that rounding cells (decreasing stretch) showed a stronger correlation between medial actin localization and apical area than did fusiform cells (increasing stretch) (**Fig. 3E**). A more notable difference was observed between junctional actin accumulation and apical area reduction, with rounding bottle cells showing much stronger correlation than the fusiform cells (**Fig. 3F**). These results indicate that medial actomyosin contractility drives apical constriction in both rounding and fusiform bottle cells, but that junctional actomyosin accumulation may be more involved in the constriction and cell shape maintenance of rounding cells than in fusiform cells.

### Knockdown of plekhg5 leads to failure in medioapical accumulation of F-actin and apical constriction

Previous studies from our groups have shown that the RhoGEF gene *plekhg5* is required for apical constriction of the bottle cells (Popov *et al*., 2018). To explore how *plekhg5* may regulate F-actin dynamics, we injected antisense splicing-blocking *plekhg5* morpholino oligo (MO) together with the RNAs of mem-mCherry and utrophin-GFP into early *Xenopus* embryos. The behaviors of F-actin in the presumptive bottle cells in the morphants were examined at early gastrula stages. Similar to cells in wild type embryos, F-actin displayed dynamic reorganization at the apical cortex. Microvilli could also be observed as distinct puncta (**Suppl. Movies 3 and 4, Fig. 4A**). However, despite the initial F-actin dynamics, progression toward enrichment of medioapical and junctional F-actin was stalled. As gastrulation proceeded, the intensity of both medial and junctional F-actin diminished over time, and apical constriction was greatly disrupted in morphants compared to wild-type cells (**Suppl. Movies 3 and 4, Fig. 4A-D**). Cell shape changes were minimal in *plekhg5* morphant cells, as morphant cells failed to form distinct round or fusiform morphology, and cell stretching was abolished (**Suppl. Movies 3 and 4**). F-actin at the cell junctions often displayed periodic intense flares before receding to the normal levels (arrowheads in **Fig. 4A**), a pattern reminiscent of junctional tear and repair (Stephenson *et al*., 2019). Taken together, the results suggest that *plekhg5* coordinates enhanced F-actin accumulation at both medial and junctional regions to efficiently reduce cell surface and impact cell shape formation.

**Figure 4.**
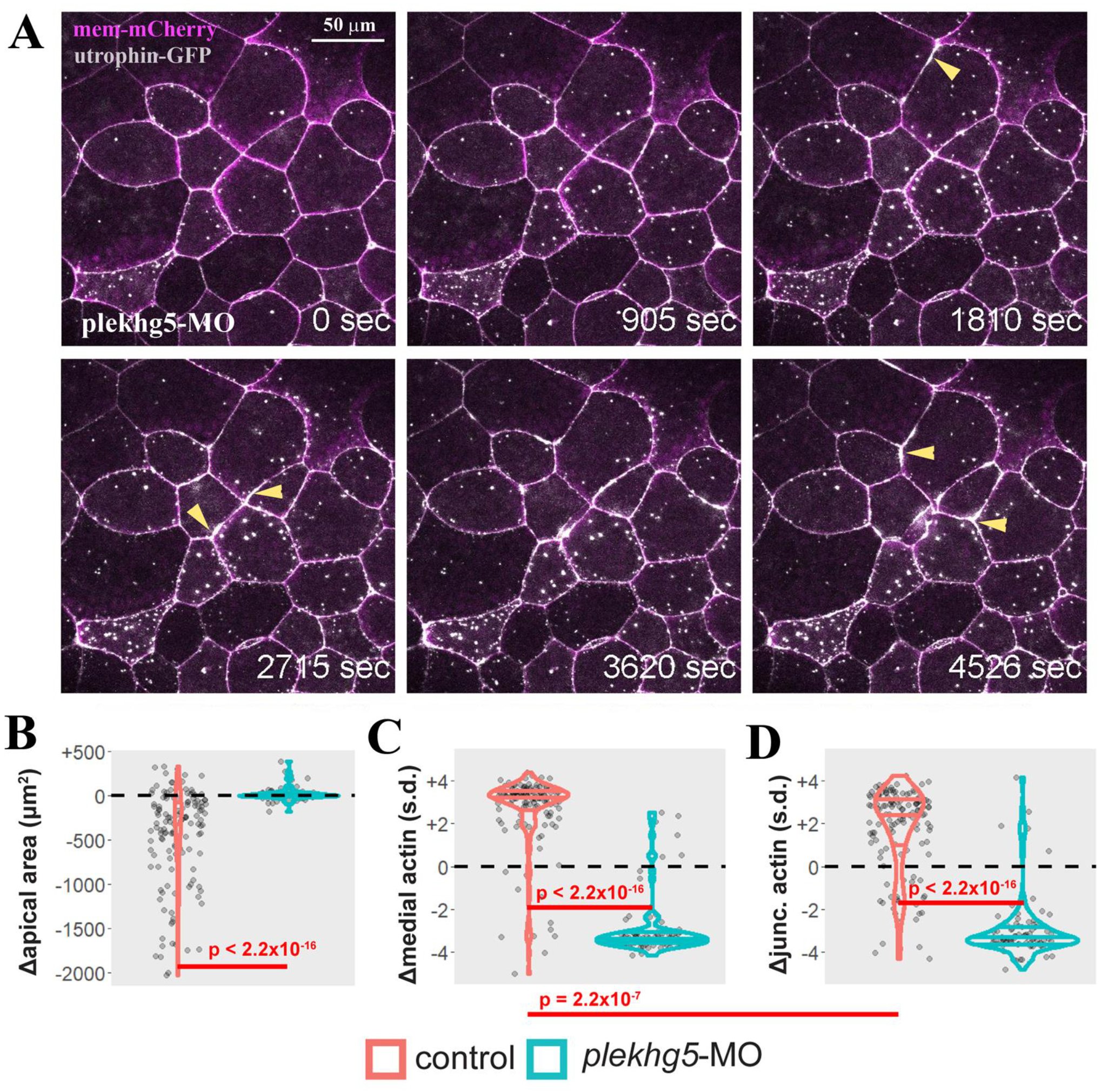
Knockdown of *plekhg5* prevents apical accumulation of F-actin and reduction of apical cell size. A) Selected frames from a time-lapse movie show that despite the initial F-actin dynamics and formation of microvilli (marked by the strong F-actin puncta within the cells), the cells with *plekhg5* knockdown fail to increase apical F-actin signal or reduce apical surface. Instead, junctional F-actin flares can be seen frequently. The arrowheads point to examples of F-actin flares. B) Quantification of apical cell area demonstrates that unlike cells in control embryos (Suppl. Movies 1 and 2), cells from *plekhg5* knockdown embryos (Suppl. Movies 3 and 4) cannot reduce their apex efficiently. C & D) While medial and junctional F-actin intensity increases during apical constriction of bottle cells in control embryos, the intensity of medial and junctional F-actin decreases in cells from *plekhg5* knockdown embryos. Each dot is an individual cell. Horizontal lines within each violin delineate quartiles along each distribution. s.d. = standard deviations. P-values were calculated via KS test.

### Coordinated action of myosin and F-actin during bottle cell formation was disrupted in plekhg5 knockdown cells

Active reduction of apical cell surface requires F-actin and non-muscle type II myosin to form the cytoskeletal contractile network. To investigate how actin and myosin coordinate during apical constriction of the bottle cells, we co-injected RNAs encoding utrophin-RFP and GFP-tagged non-muscle myosin heavy chain IIB (GFP-NMHC-IIB, or myoIIB-GFP) into the dorsal marginal zone of 4-to 8-cell stage embryos. Time-lapse microscopy revealed that like F-actin, MyoIIB was located mainly at the cell junctions prior to apical constriction. During bottle cell formation, the intensity of both F-actin and MyoIIB signals was increased, with the concurrent reduction of apical surface areas (**Suppl. Movies 5 and 6, Fig. 5**). The enhanced actomyosin signals concentrated mainly under the apical cell membrane instead of at the cell junctions. When captured at a later stage of apical constriction when bottle cells had initiated invagination, the highest actomyosin accumulation was seen to center in the medial region of the bottle cells. Both F-actin and MyoIIB formed intense signal patches in the medioapical areas. Whereas F-actin could also be detected clearly at the cell junctions, MyoIIB signals at junctions were less distinct compared to their extreme medial enrichment (**Suppl. Movie 6, Fig. 5B&C**).Apical constriction seemed to have reached its maximal extent by this stage, as cells displayed only nominal changes in apical areas while maintaining strong medioapical actomyosin signals as they invaginated further into the interior of the embryos.

**Figure 5.**
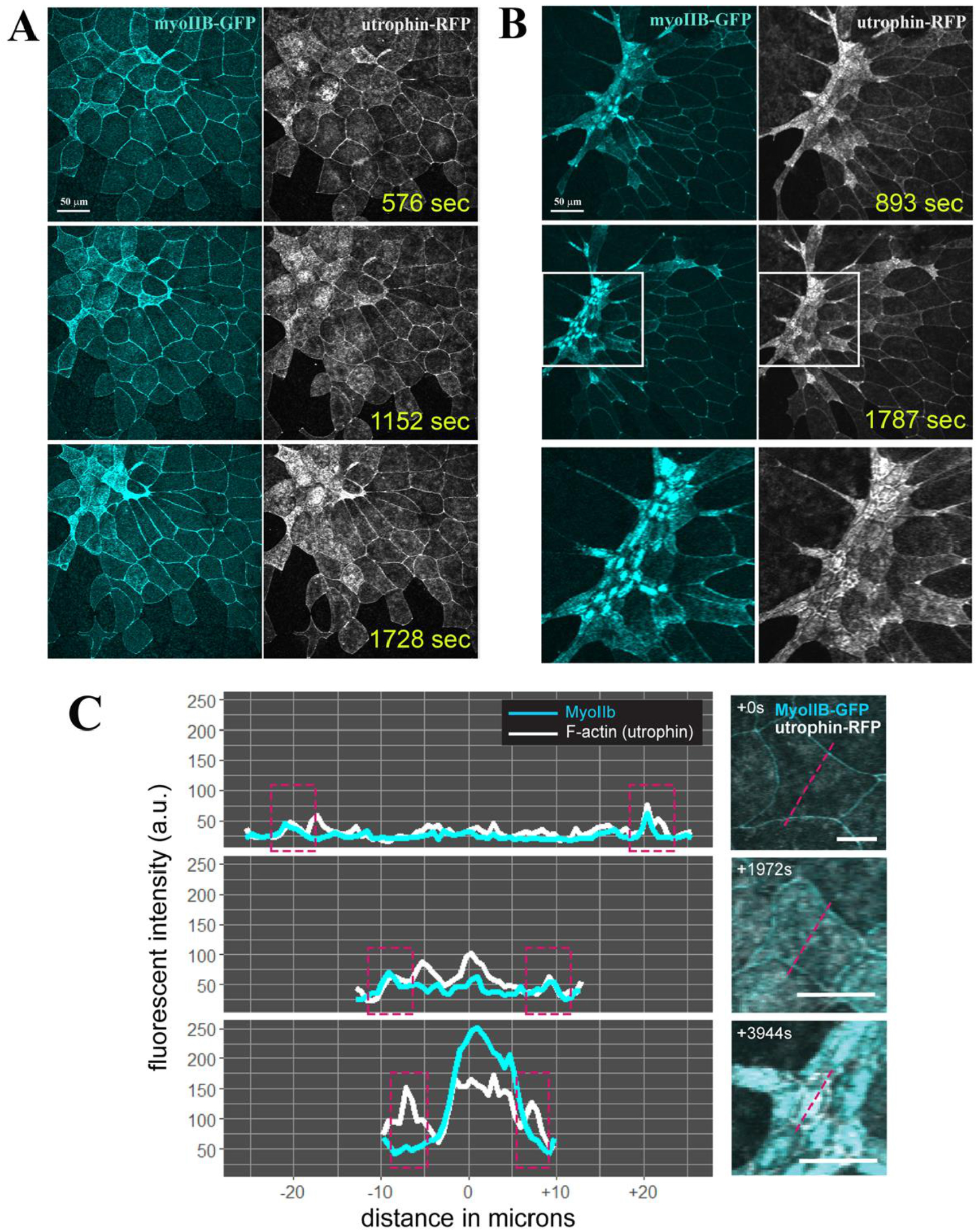
Coordinated enrichment of apical myosin IIB and F-actin during apical constriction of bottle cells. A) Selected frames of a time-lapse movie reveal coordinated increase in MyoIIB and F-actin signals in the apical cortex of the bottle cells. The signal intensity of actomyosin inversely correlates with apical cell area of the cells. The stretching of neighboring non-constricting cells can also be seen. B) During the late stage of apical constriction, MyoIIB signal is downregulated from cell junctions and concentrates in the center of the bottle cells. F-actin signal can be seen in both the cell junctions and the medioapical region. A close-up view of the boxed regions is shown in the bottom panels. C) Histogram of F-actin and MyoIIB signal intensity at the beginning, in the middle, and at the end of the apical constriction of the bottle cells reveals distinct patterns. While both medial and junctional F-actin and MyoIIB signals increase initially during bottle cell formation, junctional MyoIIB is reduced at the end of apical constriction while junctional F-actin remains strong. Approximate junctional region highlighted by dashed magenta boxes in left panels. Dashed magenta lines in right panels indicate quantified region of each cell.

To examine how *plekhg5* regulates coordination between F-actin and myosin, we tracked the dynamics of F-actin and MyoIIB simultaneously in the *plekhg5* morphant embryos (**Suppl. Movie 7, Fig. 6A&B**). As described above, F-actin displayed dynamic reorganization at the apical cell cortex and formed the punctate structures that likely reflected the formation of a limited number of microvilli in the presumptive bottle cells. MyoIIB, however, stayed mainly at the junctions of morphant cells with only a weak signal observed under the apical membrane. Medial MyoIIB accumulation was also extremely defective in *plekhg5* morphant cells (**Fig. 6C**). Junctional MyoIIB in morphant cells was not uniform and formed discrete patches that were fluid and moved along the cell-cell contacts. Over the course of imaging, *plekhg5* morphant cells failed to increase junctional MyoIIB to the same extent as control cells (**Fig. 6D**), but the major disruption was seen in the medial region where MyoIIB failed to accumulate in morphant cells (**Fig.6C&D**). Together these results indicate that Plekhg5 is primarily activating apical constriction through actomyosin recruitment and activation at the medial surfaces of bottle cells.

**Figure 6.**
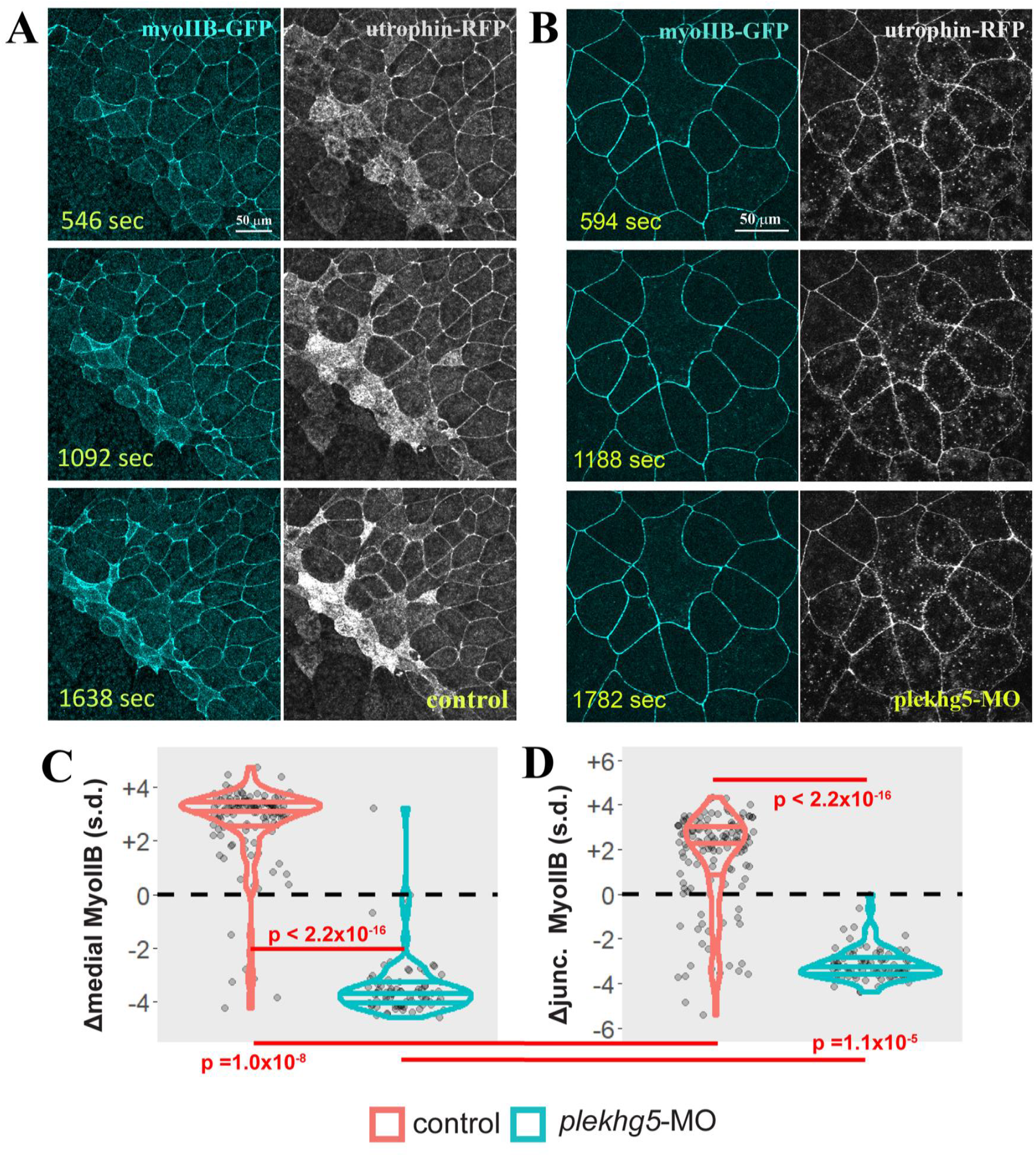
Knockdown of *plekhg5* prevents medial accumulation of MyoIIB. A&B) Unlike cells in control embryos, in *plekhg5* knockdown embryos, initial F-actin dynamics remains in the apical cortex, but MyoIIB fails to show up in the medial domain. No actomyosin enrichment is observed with progression of time. C&D) Quantification of medial and junctional MyoIIB intensity shows that unlike in control bottle cells where the intensity of MyoIIB increases, medial and junctional MyoIIB decrease in cells with *plekhg5* knockdown. Each dot is an individual cell. Horizontal lines within each violin delineate quartiles along each distribution. s.d. = standard deviations. P-values were calculated via KS test.

## Discussion

Apical constriction is a fundamental cellular process employed reiteratively to remodel epithelia during embryogenesis and in adults (Sawyer *et al*., 2010). Though actomyosin contraction is shown to be a major driving force underlying apical constriction, variations exist in terms of location, timing, and regulation of actomyosin activities as well as the relationship between actomyosin dynamics and cell shape changes during epithelial morphogenesis (Martin and Goldstein, 2014). To gain insight into molecular control of actomyosin contractile machinery during apical constriction, we used the *Xenopus* bottle cells as our model in this study and tracked simultaneously actomyosin behaviors and changes in apical cell areas both in control embryos and in embryos with knockdown of a critical RhoGEF gene *plekhg5*. Our results demonstrate heterogeneity of bottle cells and uncover an important function of *plekhg5* in organizing apical actomyosin network.

### Heterogeneous morphology of bottle cells around the blastopore lip

Bottle cells form during *Xenopus* gastrulation with drastic shrinkage of apical cell surface, striking cell elongation, and conspicuous expansion of basal cell compartment (Keller, 1981; Hardin and Keller, 1988). Previous scanning EM studies of *en face* and sagittal views of the bottle cells provide detailed morphology of these cells, and time-lapse microscopy has been used to track bottle cell formation (Keller, 1981; Hardin and Keller, 1988). However, the dynamic process leading to the cell shape changes of bottle cells has not been fully captured.

Using live imaging to track apical cell surface, we show here that two distinct groups of bottle cells emerge during gastrulation. They differ not only in morphology, with one round and the other fusiform in shape, but also in the onset of apical constriction, with the round bottle cells contracting earlier than the fusiform cells. The round, isotropically constricting cells tend to form clusters several cell diameters away and appear to actively reduce their apices autonomously. The fusiform bottle cells initiate their constriction following the round cells and seem to be influenced by their neighboring clusters of isotropically constricting cells to stretch along the circumference of the blastopore.

The heterogenous population of bottle cells are apparently required to maintain the circumference of the blastopore, as the circumference would have diminished once the bottle cells form and spread if all cells constricted in an isotropic manner. This would result in premature infolding of vegetal endoderm and impede the ordered progression of mesendoderm involution. Consistent with the geometric and mechanical constraints of the gastrulating embryos, the cells abutting the bottle cells adopt different shapes as well, with the cells on the animal-pole side to the round bottle cells displaying strong stretching along the animal-vegetal direction, whereas the cells adjacent to the fusiform bottle cells were less stretched (**Fig. 3C&D**).

### Medioapical actomyosin drives apical constriction

Our quantitative analysis of actomyosin behaviors reveals dynamics across the apical cortex during bottle cell formation. Though both junctional and medial actomyosin signals increase as apical surface area is reduced, the strongest enhancement of actomyosin intensity is seen in the medial domain. The deployment of medioapical actomyosin may provide several features consistent with bottle cell biology.

First, medial contractility could better enable cell-autonomous asynchronous contraction seen in the clusters of bottle cells some distance apart. Junctional actomyosin contractility may transmit tension to all of a cell’s neighbors during constriction, thus favoring spreading of constriction continuously around the circumference, a phenomenon inconsistent with the spaced-out clusters of cells that independently initiate constriction during blastopore lip formation (**Fig. 3A**).

Second, both endocytosis of surface membrane and generation of microvilli can reduce apical area (Keller, 1981; Hardin and Keller, 1988; Kurth and Hausen, 2000; Lee and Harland, 2010), and we have observed emerging microvilli even before large scale reduction of cell apex (**Fig. 1C**). The microvilli become especially dense as bottle cells take their mature shape. It thus appears that one main task of medioapical actomyosin is to generate and maintain high-density microvilli at the apical cell surface to facilitate apical constriction. Junctional actomyosin would theoretically be less efficient in producing cortical forces to generate dense apical microvilli. Over the period that we observed, the bottle cells invaginate slightly into the interior of the embryos, and junctional myosin is seen to be downregulated whereas the medial myosin remains exceedingly strong (**Fig. 5**). This disconnection between medial and junctional myosin may help to reduce junctional tension without relaxing apical cell area so that epithelial sheet integrity can be preserved as the bottle cells maintain their shapes during mesendoderm involution before they respread at the end of gastrulation (Keller, 1981; Hardin and Keller, 1988).

Third, the amount of medial versus junctional actomyosin activities may determine whether cells adopt round or fusiform morphology. While medial actomyosin accumulation correlates with apical area in both round and fusiform bottle cells, junctional actomyosin more strongly correlates area in cells that become rounder (**Fig. 3**). This may be due to the asymmetric junctional tension experienced by the late forming fusiform bottle cells, whereby high tension exerted by neighboring round cells dominates junctional actomyosin activities over those at the junctions non-adjacent to the round cells. The heterogeneity in junctional actomyosin in fusiform cells would negatively impact its correlation with apical area.

### Plekhg5 controls apical actomyosin accumulation to initiate apical constriction in bottle cells

The timing of concentrated actomyosin activities and the apical localization of the contractile meshwork need to be regulated tightly to allow accurate and reproducible morphogenesis. Many regulators of apical constriction have been reported in different tissue contexts and/or organisms, with the factors controlling the actions of the Rho family small GTPases among the prominent ones. The *Drosophila* DRhoGEF2 and the vertebrate GEF-H1/Arhgef2, which stimulate RhoA activation, are shown to act in the apical domain to promote actomyosin assembly and facilitate apical constriction (Barrett *et al*., 1997; Hacker and Perrimon, 1998; Nikolaidou and Barrett, 2004; Itoh *et al*., 2014). Conversely, the Cdc42 GAP protein PAC-1, which acts to inactivate Cdc42, is reported to function in the basolateral regions of *C. elegans* endodermal cells to limit Cdc42 activity only to the apical domain (Lee and Goldstein, 2003; Anderson *et al*., 2008; Chan and Nance, 2013; Marston *et al*., 2016). In *Xenopus*, we have previously identified a RhoGEF gene, *plekhg5*, that regulates apical constriction of bottle cells in RhoA-dependent manner (Popov *et al*., 2018). Plekhg5 is located in the apical domain and its activity is required for actomyosin accumulation underneath the apical membrane. However, how *plekhg5* controls actomyosin dynamics had not been described.

In this study, we show that knockdown of *plekhg5* does not affect initial F-actin dynamics in the apical cortex, and a limited number of microvilli can also form in the absence of *plekhg5*. This suggests that other regulators exist to stimulate F-actin remodeling when gastrulation starts. However, *plekhg5* is crucial for maintaining and enhancing actomyosin activities in bottle cells. Both medioapical and junctional actomyosin signals are reduced with time when *plekhg5* is knocked down. It is likely that Plekhg5 carries out the task via downstream effectors of activated RhoA, including ROCK and Diaphanous (Dia) (Narumiya *et al*., 2009). ROCK can phosphorylate regulatory myosin light chain to activate myosin contractility, whereas Dia is a formin-domain-containing protein that can promote actin bundle formation. Apically localized Plekhg5 can therefore stimulate local assembly and contraction of actomyosin to facilitate reduction of apical surface area. It is interesting to note that while *plekhg5* is expressed in cells around the blastopore lip, not all the blastopore lip cells express *plekhg5* at the same time. Instead, the gene is expressed in a salt-and-pepper fashion in subsets of the cells in the lip region (**Suppl. Fig. 1**). Considering that we observe the round and the fusiform bottle cells that differ in temporal onset and possibly autonomy in force generation, we propose that *plekhg5*-expressing cells may initiate isotropic apical constriction in discrete cells that contribute to early forming round bottle cells, whereas the *plekhg5* non-expressing cells constrict later and take the fusiform shape. Knockdown of *plekhg5* would prevent formation of both types of bottle cells, as fusiform cells seem to depend on the appearance of round cells. Though this model is consistent with our data, we do not have direct evidence currently.

### Plekhg5 and Shroom3 are mechanistically distinct activators of apical constriction during Xenopus embryogenesis

When compared with apical constriction in the context of neural tube closure in *Xenopus* (Baldwin *et al*., 2022), we find both common and unique features. In both bottle cells and cells within the anterior neural plate, medioapical actomyosin appears to be the main driving force for apical constriction. However, the mechanisms of actomyosin regulation seem to differ between these tissues. As we have shown previously (Baldwin *et al*., 2022), the actin-binding protein Shroom3 is a critical regulator of apical constriction in neural ectoderm cells. Shroom3 controls N-cadherin localization and coupling of actin contraction to effectively reduce apical surface area. Mutation of *shroom3* does not entirely prevent medial actin accumulation but precludes actin-driven reduction of apical surface area. In comparison, none of the *shroom* family genes are expressed in bottle cells (Lee *et al*., 2009). Apical constriction of bottle cells therefore proceeds independent of *shroom* function and relies on the activity of the RhoGEF gene *plekhg5*.

Unlike in *shroom3* depleted cells, actin fails to accumulate either in medial or junctional regions in *plekhg5* knockdown cells. This suggests that *plekhg5* plays a more fundamental role than *shroom3* in assembly of apical actomyosin network. As both Shroom3 and Plekhg5 are apically localized and can activate ROCK via either direct binding to ROCK or Rho activation (Ngok and Anastasiadis, 2013; Das *et al*., 2014; Mack and Georgiou, 2014), the difference in their function in regulating apical actin accumulation may reflect other distinct partners and downstream effectors of these two factors. Identification of molecular components involved in control of apical constriction by Shroom3 and Plekhg5 can provide valuable information on distinct mechanisms regulating an important cellular process underlying epithelial morphogenesis.

In summary, we show in this study that bottle cells are not homogenous and take different shapes. Formation of bottle cells rely more on medioapical rather than on junctional actomyosin meshwork, and the RhoGEF protein Plekhg5 is essential for enhancing actomyosin activity so that apical constriction can proceed to generate epithelial deformation during *Xenopus* gastrulation. Though the medioapical actomyosin meshwork seems to be a shared feature of apical constriction with many other species, microvilli appear to be more specific for reducing the large surface area in the bottle cells of *Xenopus*. It will be interesting to investigate in future whether fusiform bottle cells constrict actively or passively, how remodeling of junctional adhesion complexes contributes to reduction of apical surface area, what functions of Rho effectors ROCK and Dia play during bottle cell formation, and how Plekhg5 expression and activity are regulated by other signals involved in cell fate determination and morphogenesis.

## Materials and Methods

### Embryo culture and injection

*Xenopus laevis* frogs were used under the institutional IACUC protocol 21371 at the University of Alabama at Birmingham. The embryos were obtained by *in vitro* fertilization. The GFP-MyoIIB construct was obtained from Addgene (Addgene plasmid #11348) (Wei and Adelstein, 2000) and cloned into the pCS105 vector. The membrane mCherry and utrophin-GFP/RFP plasmids were used as described before (Burkel *et al*., 2007). RNAs were synthesized from the linearized plasmids using the *in vitro* mMessage mMachine RNA synthesis kit (Ambion). 100-250pg of Mem-mCherry and utrophin-GFP/RFP RNAs and 1-2ng myoIIB-GFP RNAs were injected into the marginal zone regions of two dorsal blastomeres of 4-to 8-cell stage embryos. The embryos were imaged around the time just before and immediately after the dorsal lip appears, from the late blastula to early gastrula stages. Half the injected embryos were cultured to the tailbud to tadpole stages to ensure that the injected RNAs did not affect the embryo development. For *plekhg5* knockdown, a previously tested antisense splicing-blocking morpholino oligo (Popov *et al*., 2018) was used at total 25 ng to 33 ng into the dorsal marginal zone of 4-cell stage embryos.

### Imaging of actomyosin dynamics in the bottle cells

The injected embryos were collected at late blastula stages, with their vitelline membrane removed, and imaged with either upright Olympus Fluoview F1000 or inverted Nikon A1R confocal microscopes. To prevent embryos from sticking to the cover glasses or imaging dishes, the glasses and the dishes were treated with 1% BSA. The embryos were positioned with the presumptive dorsal lip region in the imaging view. Time-lapse microscopy was performed with z-stack to cover the entire curved surface in view and no pause in between time points (continuous run). This would end up with images with 4 to 17 z-stacks and a time interval of about 16 to 22 seconds. Maximum intensity projection was carried out using FIJI and the resulting movies were shown in the Supplemental Materials.

### Quantitative image analysis

Cells were segmented and tracked using EPySeg and then EPySeg outputs were manually corrected using Tissue Analyzer (Aigouy *et al*., 2010; Aigouy *et al*., 2016; Aigouy *et al*., 2020). Tissue Analyzer was used to break the apical surfaces of cells into two domains: a “medial” domain which encompasses the apical surface within the cell-cell junctions, and a “junctional” domain which is composed solely of the cell-cell junctions (**Figure 2B**). Tissue Analyzer and FIJI (Schindelin *et al*., 2012) were further used to calculate mean fluorescent intensities and physical size parameters at each of these domains. Tissue Analyzer databases were imported to R and further analyzed and manipulated primarily using the tidyverse package (Wickham *et al*., 2019; R Core Team, 2020). Frame-intervals for each movie varied slightly, ranging from 16 to 22 seconds, so parameters within each cell were averaged over a 7-frame window to account for “noise” within our measurements. To account for differences in initial size and intensity of fluorescent markers, parameters were standardized by mean-centering the data for each individual cell track to zero and then dividing the resulting mean-centered values by the standard deviation of each track. Thus, parameters are displayed both in standard deviations as well as square microns or arbitrary units, allowing for relative dynamics to be compared between cells in addition to raw changes in parameters (Baldwin *et al*., 2022). Kolmogorov-Smirnov tests were performed using the *ks*.*test* function in R.

## Acknowledgements

The work is supported by the NIH grant R01 GM127371 (CC and RK), R01HD099191 (JBW), and NIH fellowship F32 HD094521 (ATB).

---

**Supplementary Figure 1.**
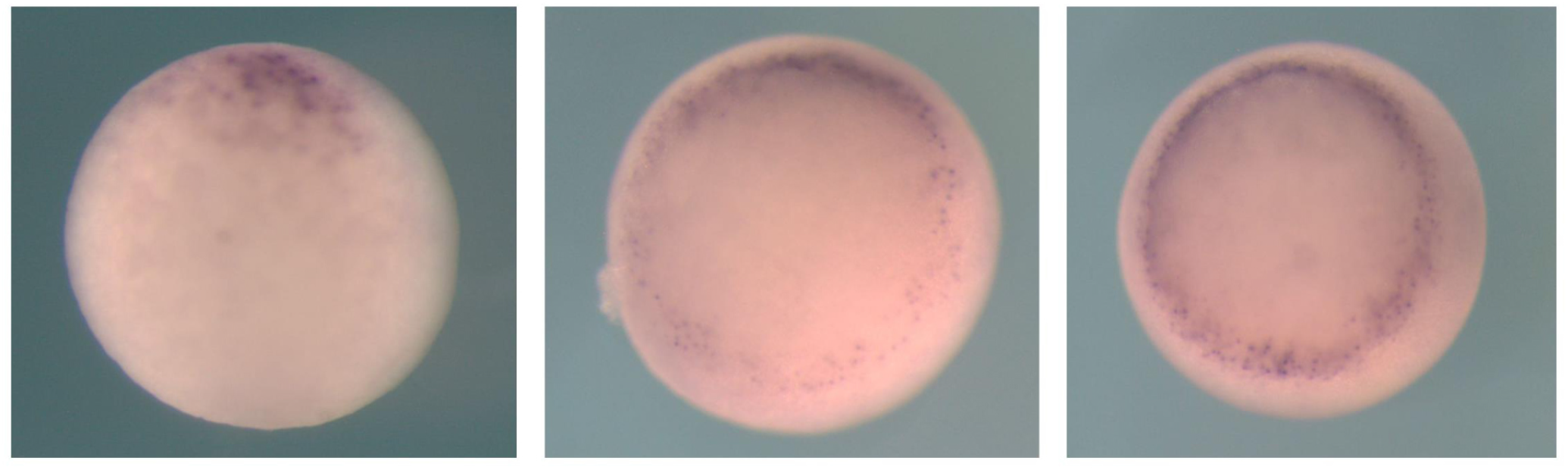
plekhg5 is expressed in a salt-and-pepper fashion in cells within the blastopore lip.

## References

Aigouy, B., Cortes, C., Liu, S., and Prud’Homme, B. (2020). EPySeg: a coding-free solution for automated segmentation of epithelia using deep learning. Development 147.

Aigouy, B., Farhadifar, R., Staple, D.B., Sagner, A., Roper, J.C., Julicher, F., and Eaton, S. (2010). Cell flow reorients the axis of planar polarity in the wing epithelium of Drosophila. Cell 142, 773–786.

Aigouy, B., Umetsu, D., and Eaton, S. (2016). Segmentation and Quantitative Analysis of Epithelial Tissues. Methods Mol Biol 1478, 227–239.

Anderson, D.C., Gill, J.S., Cinalli, R.M., and Nance, J. (2008). Polarization of the C. elegans embryo by RhoGAP-mediated exclusion of PAR-6 from cell contacts. Science 320, 1771–1774.

Baldwin, A.T., Kim, J., Seo, H., and Wallingford, J.B. (2022). Global analysis of cell behavior and protein localization dynamics reveals region-specific functions for Shroom3 and N-cadherin during neural tube closure. Elife (Cambridge) 11, e66704.

Barrett, K., Leptin, M., and Settleman, J. (1997). The Rho GTPase and a putative RhoGEF mediate a signaling pathway for the cell shape changes in Drosophila gastrulation. Cell 91, 905–915.

Bement, W.M., Forscher, P., and Mooseker, M.S. (1993). A novel cytoskeletal structure involved in purse string wound closure and cell polarity maintenance. J Cell Biol 121, 565–578.

Booth, A.J.R., Blanchard, G.B., Adams, R.J., and Roper, K. (2014). A dynamic microtubule cytoskeleton directs medial actomyosin function during tube formation. Dev Cell 29, 562–576.

Burkel, B.M., von Dassow, G., and Bement, W.M. (2007). Versatile fluorescent probes for actin filaments based on the actin-binding domain of utrophin. Cell Motil Cytoskeleton 64, 822–832.

Chan, E., and Nance, J. (2013). Mechanisms of CDC-42 activation during contact-induced cell polarization. J Cell Sci 126, 1692–1702.

Chanet, S., Miller, C.J., Vaishnav, E.D., Ermentrout, B., Davidson, L.A., and Martin, A.C. (2017). Actomyosin meshwork mechanosensing enables tissue shape to orient cell force. Nat Commun 8, 15014.

Choi, S.C., and Sokol, S.Y. (2009). The involvement of lethal giant larvae and Wnt signaling in bottle cell formation in Xenopus embryos. Dev Biol 336, 68–75.

Christodoulou, N., and Skourides, P.A. (2015). Cell-Autonomous Ca(2+) Flashes Elicit Pulsed Contractions of an Apical Actin Network to Drive Apical Constriction during Neural Tube Closure. Cell Rep 13, 2189–2202.

Das, D., Zalewski, J.K., Mohan, S., Plageman, T.F., VanDemark, A.P., and Hildebrand, J.D. (2014). The interaction between Shroom3 and Rho-kinase is required for neural tube morphogenesis in mice. Biol Open 3, 850–860.

Goldstein, B., and Nance, J. (2020). Caenorhabditis elegans Gastrulation: A Model for Understanding How Cells Polarize, Change Shape, and Journey Toward the Center of an Embryo. Genetics 214, 265–277.

Hacker, U., and Perrimon, N. (1998). DRhoGEF2 encodes a member of the Dbl family of oncogenes and controls cell shape changes during gastrulation in Drosophila. Genes Dev 12, 274–284.

Hardin, J., and Keller, R. (1988). The behaviour and function of bottle cells during gastrulation of Xenopus laevis. Development 103, 211–230.

Itoh, K., Ossipova, O., and Sokol, S.Y. (2014). GEF-H1 functions in apical constriction and cell intercalations and is essential for vertebrate neural tube closure. J Cell Sci 127, 2542–2553.

Jodoin, J.N., Coravos, J.S., Chanet, S., Vasquez, C.G., Tworoger, M., Kingston, E.R., Perkins, L.A., Perrimon, N., and Martin, A.C. (2015). Stable Force Balance between Epithelial Cells Arises from F-Actin Turnover. Dev Cell 35, 685–697.

Kamran, Z., Zellner, K., Kyriazes, H., Kraus, C.M., Reynier, J.B., and Malamy, J.E. (2017). In vivo imaging of epithelial wound healing in the cnidarian Clytia hemisphaerica demonstrates early evolution of purse string and cell crawling closure mechanisms. BMC Dev Biol 17, 17.

Keller, R.E. (1981). An experimental analysis of the role of bottle cells and the deep marginal zone in gastrulation of Xenopus laevis. J Exp Zool 216, 81–101.

Kiehart, D.P., Crawford, J.M., Aristotelous, A., Venakides, S., and Edwards, G.S. (2017). Cell Sheet Morphogenesis: Dorsal Closure in Drosophila melanogaster as a Model System. Annu Rev Cell Dev Biol 33, 169–202.

Ko, C.S., Tserunyan, V., and Martin, A.C. (2019). Microtubules promote intercellular contractile force transmission during tissue folding. J Cell Biol 218, 2726–2742.

Kurth, T. (2005). A cell cycle arrest is necessary for bottle cell formation in the early Xenopus gastrula: integrating cell shape change, local mitotic control and mesodermal patterning. Mech Dev 122, 1251–1265.

Kurth, T., and Hausen, P. (2000). Bottle cell formation in relation to mesodermal patterning in the Xenopus embryo. Mech Dev 97, 117–131.

Lang, R.A., Herman, K., Reynolds, A.B., Hildebrand, J.D., and Plageman, T.F., Jr. (2014). p120-catenin-dependent junctional recruitment of Shroom3 is required for apical constriction during lens pit morphogenesis. Development 141, 3177–3187.

Le, T.P., and Chung, S. (2021). Regulation of apical constriction via microtubule- and Rab11-dependent apical transport during tissue invagination. Mol Biol Cell 32, 1033–1047.

Lee, C., Le, M.P., and Wallingford, J.B. (2009). The shroom family proteins play broad roles in the morphogenesis of thickened epithelial sheets. Dev Dyn 238, 1480–1491.

Lee, C., Scherr, H.M., and Wallingford, J.B. (2007). Shroom family proteins regulate gamma-tubulin distribution and microtubule architecture during epithelial cell shape change. Development 134, 1431–1441.

Lee, J.Y., and Goldstein, B. (2003). Mechanisms of cell positioning during C. elegans gastrulation. Development 130, 307–320.

Lee, J.Y., and Harland, R.M. (2007). Actomyosin contractility and microtubules drive apical constriction in Xenopus bottle cells. Dev Biol 311, 40–52.

Lee, J.Y., and Harland, R.M. (2010). Endocytosis is required for efficient apical constriction during Xenopus gastrulation. Curr Biol 20, 253–258.

Mack, N.A., and Georgiou, M. (2014). The interdependence of the Rho GTPases and apicobasal cell polarity. Small GTPases 5, 10.

Marston, D.J., Higgins, C.D., Peters, K.A., Cupp, T.D., Dickinson, D.J., Pani, A.M., Moore, R.P., Cox, A.H., Kiehart, D.P., and Goldstein, B. (2016). MRCK-1 Drives Apical Constriction in C. elegans by Linking Developmental Patterning to Force Generation. Curr Biol 26, 2079–2089.

Martin, A.C., and Goldstein, B. (2014). Apical constriction: themes and variations on a cellular mechanism driving morphogenesis. Development 141, 1987–1998.

Martin, A.C., Kaschube, M., and Wieschaus, E.F. (2009). Pulsed contractions of an actin-myosin network drive apical constriction. Nature 457, 495–499.

Mason, F.M., Tworoger, M., and Martin, A.C. (2013). Apical domain polarization localizes actin-myosin activity to drive ratchet-like apical constriction. Nat Cell Biol 15, 926–936.

Massarwa, R., Schejter, E.D., and Shilo, B.Z. (2009). Apical secretion in epithelial tubes of the Drosophila embryo is directed by the Formin-family protein Diaphanous. Dev Cell 16, 877–888.

Mulinari, S., Barmchi, M.P., and Hacker, U. (2008). DRhoGEF2 and diaphanous regulate contractile force during segmental groove morphogenesis in the Drosophila embryo. Mol Biol Cell 19, 1883–1892.

Nakajima, H., and Tanoue, T. (2011). Lulu2 regulates the circumferential actomyosin tensile system in epithelial cells through p114RhoGEF. J Cell Biol 195, 245–261.

Nakajima, H., and Tanoue, T. (2012). The circumferential actomyosin belt in epithelial cells is regulated by the Lulu2-p114RhoGEF system. Small GTPases 3, 91–96.

Narumiya, S., Tanji, M., and Ishizaki, T. (2009). Rho signaling, ROCK and mDia1, in transformation, metastasis and invasion. Cancer Metastasis Rev 28, 65–76.

Ngok, S.P., and Anastasiadis, P.Z. (2013). Rho GEFs in endothelial junctions: Effector selectivity and signaling integration determine junctional response. Tissue Barriers 1, e27132.

Nikolaidou, K.K., and Barrett, K. (2004). A Rho GTPase signaling pathway is used reiteratively in epithelial folding and potentially selects the outcome of Rho activation. Curr Biol 14, 1822–1826.

Ossipova, O., Chuykin, I., Chu, C.W., and Sokol, S.Y. (2015). Vangl2 cooperates with Rab11 and Myosin V to regulate apical constriction during vertebrate gastrulation. Development 142, 99–107.

Plageman, T.F., Jr., Chauhan, B.K., Yang, C., Jaudon, F., Shang, X., Zheng, Y., Lou, M., Debant, A., Hildebrand, J.D., and Lang, R.A. (2011). A Trio-RhoA-Shroom3 pathway is required for apical constriction and epithelial invagination. Development 138, 5177–5188.

Popov, I.K., Ray, H.J., Skoglund, P., Keller, R., and Chang, C. (2018). The RhoGEF protein Plekhg5 regulates apical constriction of bottle cells during gastrulation. Development 145.

R Core Team. (2020). R: A Language and Environment for Statistical Computing, Vienna, Austria: R Foundation for Statistical Computing.

Rogers, S.L., Wiedemann, U., Hacker, U., Turck, C., and Vale, R.D. (2004). Drosophila RhoGEF2 associates with microtubule plus ends in an EB1-dependent manner. Curr Biol 14, 1827–1833.

Roh-Johnson, M., Shemer, G., Higgins, C.D., McClellan, J.H., Werts, A.D., Tulu, U.S., Gao, L., Betzig, E., Kiehart, D.P., and Goldstein, B. (2012). Triggering a cell shape change by exploiting preexisting actomyosin contractions. Science 335, 1232–1235.

Rousso, T., Shewan, A.M., Mostov, K.E., Schejter, E.D., and Shilo, B.Z. (2013). Apical targeting of the formin Diaphanous in Drosophila tubular epithelia. Elife 2, e00666.

Sawyer, J.M., Harrell, J.R., Shemer, G., Sullivan-Brown, J., Roh-Johnson, M., and Goldstein, B. (2010). Apical constriction: a cell shape change that can drive morphogenesis. Dev Biol 341, 5–19.

Schindelin, J., Arganda-Carreras, I., Frise, E., Kaynig, V., Longair, M., Pietzsch, T., Preibisch, S., Rueden, C., Saalfeld, S., Schmid, B., Tinevez, J.Y., White, D.J., Hartenstein, V., Eliceiri, K., Tomancak, P., and Cardona, A. (2012). Fiji: an open-source platform for biological-image analysis. Nat Methods 9, 676–682.

Stephenson, R.E., Higashi, T., Erofeev, I.S., Arnold, T.R., Leda, M., Goryachev, A.B., and Miller, A.L. (2019). Rho Flares Repair Local Tight Junction Leaks. Dev Cell 48, 445–459 e445.

Wei, Q., and Adelstein, R.S. (2000). Conditional expression of a truncated fragment of nonmuscle myosin II-A alters cell shape but not cytokinesis in HeLa cells. Mol Biol Cell 11, 3617–3627.

Wickham, H., Averick, M., Bryan, J., Chang, W., D’Agostino McGowan, L., François, R., Grolemund, G., Hayes, A., Henry, L., Hester, J., Kuhn, M., Pedersen, T.L., Miller, E., Bache, S.M., Müller, K., Robinson, D., Seidel, D.P., Spinu, V., Takahashi, K., Vaughan, D., Wilke, C., Woo, K., and Yutani, H. (2019). Welcome to the Tidyverse. Journal of Open Source Software.

Yano, T., Tsukita, K., Kanoh, H., Nakayama, S., Kashihara, H., Mizuno, T., Tanaka, H., Matsui, T., Goto, Y., Komatsubara, A., Aoki, K., Takahashi, R., Tamura, A., and Tsukita, S. (2021). A microtubule-LUZP1 association around tight junction promotes epithelial cell apical constriction. EMBO J 40, e104712.

